# Genetic drift and purifying selection shape within-host influenza A virus populations during natural swine infections

**DOI:** 10.1101/2023.10.23.563581

**Authors:** David VanInsberghe, Dillon S. McBride, Juliana DaSilva, Thomas J. Stark, Max S.Y. Lau, Samuel S. Shepard, John R. Barnes, Andrew S. Bowman, Anice C. Lowen, Katia Koelle

## Abstract

Patterns of within-host influenza A virus (IAV) diversity and evolution have been described in natural human infections, but these patterns remain poorly characterized in non-human hosts. Elucidating these dynamics is important to better understand IAV biology and the evolutionary processes that govern spillover into humans. Here, we sampled an IAV outbreak in pigs during a week-long county fair to characterize viral diversity and evolution in this important reservoir host. Nasal wipes were collected on a daily basis from all pigs present at the fair, yielding up to 421 samples per day. Subtyping of PCR-positive samples revealed the co-circulation of H1N1 and H3N2 subtype IAVs. PCR-positive samples with robust Ct values were deep-sequenced, yielding 506 sequenced samples from a total of 253 pigs. Based on higher-depth re-sequenced data from a subset of these initially sequenced samples (260 samples from 168 pigs), we characterized patterns of within-host IAV genetic diversity and evolution. We find that IAV genetic diversity in single-subtype infected pigs is low, with the majority of intra-host single nucleotide variants (iSNVs) present at frequencies of <10%. The ratio of the number of nonsynonymous to the number of synonymous iSNVs is significantly lower than under the neutral expectation, indicating that purifying selection shapes patterns of within-host viral diversity in swine. The dynamic turnover of iSNVs and their pronounced frequency changes further indicate that genetic drift also plays an important role in shaping IAV populations within pigs. Taken together, our results highlight similarities in patterns of IAV genetic diversity and evolution between humans and swine, including the role of stochastic processes in shaping within-host IAV dynamics.

## Introduction

Pigs host a rich and epidemiologically important reservoir of influenza A virus (IAV) genetic diversity [1,2]. The swine IAV gene pool is strongly shaped by frequent spill-over between pigs and humans [3–5] and occasional transmission of avian IAV to pigs [6]. There are three main subtypes circulating within swine populations: H1N1, H1N2, and H3N2 [7]. Each subtype contains multiple clades that are antigenically distinct, including those that evolved following reverse zoonosis of human seasonal IAV [8]. The evolutionary dynamics of these IAV lineages are an ongoing concern for the swine industry because infections substantially impact animal welfare and productivity. They also pose a significant risk for global public health, as was laid bare with the 2009 H1N1 pandemic. This pandemic was caused by a reassortant swine virus carrying gene segments derived from multiple swine, human, and avian IAV lineages [9,10]. Since that event, hundreds of self-limiting swine-to-human zoonoses have been documented in the USA, a large proportion of which occur at agricultural fairs [11,12]. These recurring transmission events highlight the importance of the swine-human interface as a source of IAV pandemics and, in turn, give a strong motivation for research into the viral evolutionary processes playing out in pigs.

Our understanding of IAV diversity in pigs at a population level has improved substantially since 2009. Increased investment in swine IAV surveillance and expanded use of genome sequencing has provided novel insight into how swine IAV circulates and evolves over time and between geographic locations [1,7,13–15]. However, the processes shaping viral evolution within individual pigs has, to date, garnered less attention. Because novel IAV variants and lineages circulating at a population level must originate within individuals hosts, knowledge of the drivers of viral evolution at the within-host scale is essential to better understand viral evolution across all scales of biological organization.

Many features of the within-host evolutionary dynamics of natural IAV infections in humans have been examined [16–20], together revealing that transmission bottlenecks are small, that within-host levels of genetic diversity are low, and that genetic drift and purifying selection are the primary drivers of within-host viral evolution. However, whether within-host IAV evolution occurs in a similar fashion in non-human hosts, in particular swine, remains unclear. Features of natural swine hosts and their management in agricultural settings could give rise to important differences. For example, pigs raised for pork typically reach only six months of age before harvest, reducing the potential for repeated IAV infection and therefore potentially limiting selection imposed by memory immune responses. Animal feeding operations may further allow for closer contact between hosts, which could result in less stringent transmission bottlenecks and therewith the potential for higher levels of IAV genetic diversity to be transmitted. Infected pigs might therefore harbor more viral genetic and phenotypic variation for selection to act upon.

Here we aim to uncover the dominant evolutionary processes shaping IAV populations within naturally infected swine through in-depth analysis of viral samples taken during the course of an IAV outbreak at a week-long agricultural fair. Dense longitudinal sampling at this fair resulted in the identification of 367 (out of 422 sampled) pigs that were infected with IAV over the course of the fair. Samples with robust Ct values were deep-sequenced, yielding a total of 506 sequenced samples. To ensure our conclusions were robust to the presence of spurious nucleotide variants, we selected a subset of these samples to be re-sequenced to a much greater depth. We characterized patterns of IAV genetic diversity in these samples and investigated the dynamics of minority variants in the subset of pigs with sequenced longitudinal samples. Based on these analyses, we evaluated the contributions of selection and genetic drift in driving the within-host evolutionary dynamics of IAV in pigs.

## Results

### Two IAV subtypes co-circulated at the fair

The agricultural fair took place in 2014 and involved the showing of a total of 423 exhibition swine. Nasal wipes were collected from all pigs present at the fair on day 1 through day 6. All 2159 nasal wipes were screened for IAV using a real-time reverse transcription-polymerase chain reaction (rRT-PCR) assay targeting the M segment. On day 1 of the fair, few pigs were found to be PCR-positive for IAV (33 of 419 samples; **Table S1**). Over the next five days, PCR positivity rates increased from 7.9% (day 1) to 88.0% (day 6; 139 of 158 samples) (**Figure 1A**). A subset of the positive samples were then tested using an IAV subtype-specific assay, which revealed that both H3N2 and H1N1 viruses were circulating at the fair (**Table S1**). Samples with an M segment rRT-PCR cycle threshold (Ct) value of ≤31.0 were selected for next generation sequencing, resulting in sequenced samples from a total of 253 pigs (**Figure 1B**; **Table S2**). These sequenced samples had a read depth across the genome of approximately 4000 reads per nucleotide (**Figure S1**).

**Figure 1.**
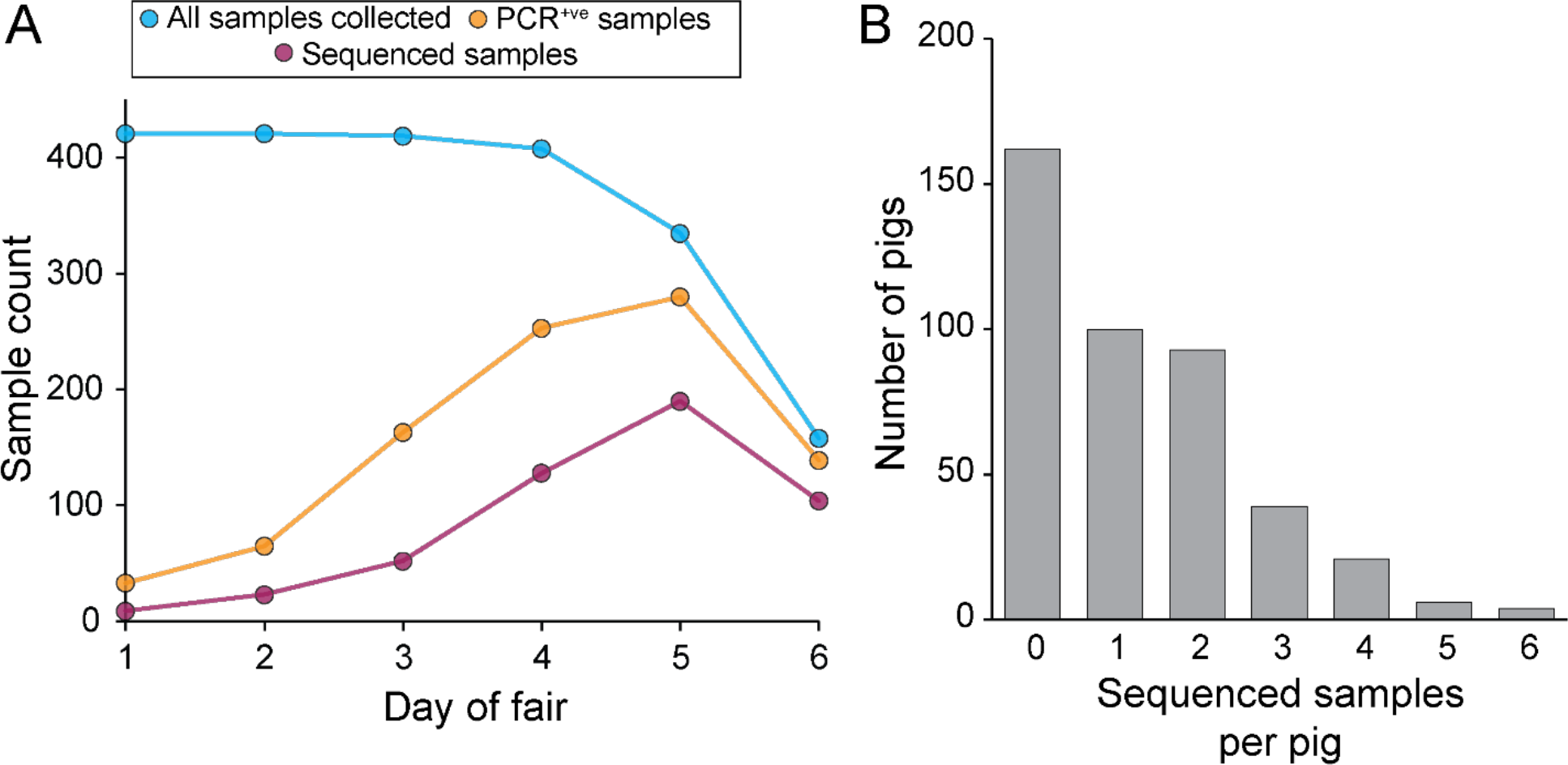
Overview of sampling and sequencing of pigs at the county fair. **(A)** Sampling, testing, and sequencing effort over the course of the fair. The decrease in the number of samples collected on days 5 and 6 stems from the early departure of a subset of the pigs. (B) Distribution of the number of sequenced samples per pig. Pigs with no sequenced samples either never tested positive for IAV or tested positive at some point but positive samples all had Ct values exceeding 31.0.

To characterize diversity of IAV genotypes present, we inferred consensus sequences for each sample for each of the eight IAV gene segments. Haplotype networks reconstructed from these consensus sequences indicated that two distinct lineages co-circulated at the fair (**Figure 2**), consistent with the subtyping results. We labeled these two lineages I and II, each comprising a set of eight gene segments that were grouped together based on the associations that were evident in the data. By comparing the outcomes of the subtype-specific tests against the assigned lineages of the consensus sequences, we were able to conclude that the hemagglutinin (HA) and neuraminidase (NA) gene segments of lineage I belonged to IAV subtype H1N1 and that the HA and NA gene segments of lineage II belonged to IAV subtype H3N2. Each of the two lineages comprised a single dominant consensus sequence and many consensus ‘singletons’. This indicates that the outbreak at the fair likely originated from a very small number of pigs (possibly only 2) that arrived at the fair already infected.

**Figure 2.**
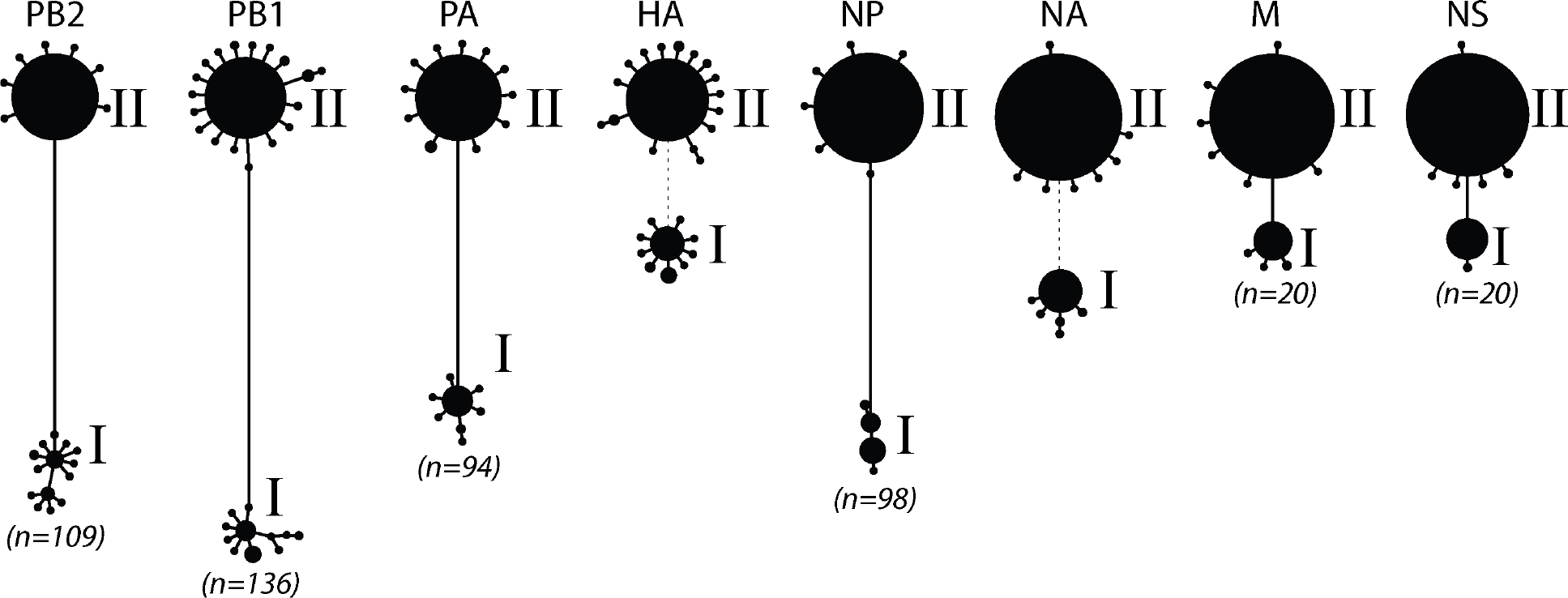
Haplotype networks of IAV segments support co-circulation of two distinct lineages at the county fair. Consensus sequences from all successfully sequenced samples were inferred and used to construct haplotype networks. Each circle represents a unique haplotype (excluding differences caused by ambiguous bases), with circle size of a haplotype being proportional to the number of consensus sequences that belong to it. Numbers in parentheses refer to the number of differentiating single nucleotide polymorphisms (dSNPs) that distinguish the dominant haplotypes from each lineage in each segment. Because of the high genetic divergence between the HA of lineage I and the HA of lineage II, and between the NA of lineage I and the NA of lineage II, dotted lines rather than solid lines are shown between the lineages.

Despite the majority of samples having consensus sequences that belonged either exclusively to lineage I or to lineage II, a small number of samples had consensus sequences with at least one gene segment belonging to a different lineage than the others (e.g., vial 14SW5957). These putatively reassortant samples indicate that at least some of the infected pigs at the fair were likely to have been heterosubtypically coinfected. To identify samples that were heterosubtypically coinfected, we looked at viral diversity below the consensus level. For this purpose, we selected representative consensus sequences of each segment from both lineages to serve as references for read mapping. We first assessed whether there was evidence for HA coinfection or NA coinfection. Because the consensus sequences of the HA and of the NA segments of lineages I and II are sufficiently distinct at the nucleotide level to support unambiguous mapping, we detected co-occurrence of these segments by mapping the samples’ reads to H1, N1, H3, and N2 reference genomes. Samples with reads mapping to only one HA and one NA subtype were classified as singly infected at each of these gene segments (**Figure 3A**; **Table S2**). Samples that had an appreciable number of reads mapping to both H3 and H1 were classified as coinfected on the HA gene segment. Similarly, samples that had an appreciable number of reads mapping to both N2 and N1 were classified as coinfected on the NA gene segment.

**Figure 3.**
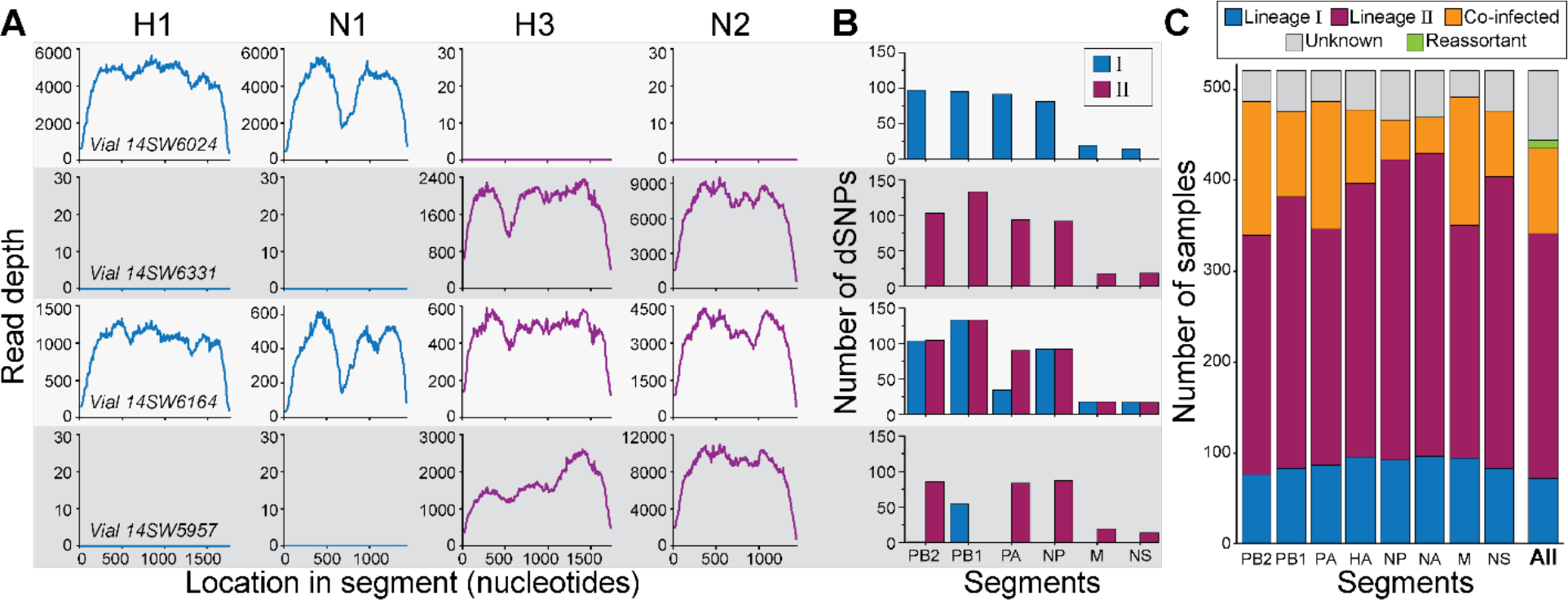
Classification of gene segments and samples into lineages. **(A)** Mapping of sample reads on to reference H1, N1, H3, and N2 gene segments for four selected samples. Based on this mapping, a sample’s HA gene segment was classified as belonging to either lineage I (with reads mapped to H1; vial#14SW6024), lineage II (with reads mapped to H3; vial#14SW6331 and vial#14SW5957), or coinfected (with reads mapped to both H1 and H3; vial#14SW6164). Similarly, a sample’s NA gene segment was classified as belonging to either lineage I (with reads mapped to N1; vial#14SW6024), lineage II (with reads mapped to N2; vial#14SW6331 and vial#14SW5957), or coinfected (with reads mapped to both N1 and N2; vial#14SW6164). (B) Detection of dSNPs characteristic of lineages I and II for each of the six internal gene segments of the four selected samples. (C) The proportion of samples classified as lineage I, II, or coinfected, by gene segment. Segments were labeled as unknown when there was insufficient coverage across the focal gene segment to determine nucleotide identities at the dSNP sites or to robustly map to the HA and NA gene segments. Samples (‘All’) were classified as belonging to lineage I, lineage II, coinfected, or reassortant based on their constituent segments. A sample was defined as a lineage I or II sample if at least five of its gene segments were successfully classified and found to be either singly infected lineage I or singly infected lineage II, with no evidence of coinfection. A sample was considered a reassortant if at least five segments were successfully classified and a combination of lineage I singly infected and lineage II singly infected segments were present in the same sample. A sample was considered coinfected if at least one of its gene segments was classified as coinfected. In panels (A) and (B), each row corresponds to a different sample classification. Vial#14SW6024 corresponds to a sample with a lineage I virus, vial#14SW6331 corresponds to a sample with a lineage II virus, vial#14SW6164 corresponds to a coinfected sample, and vial#14SW5957 corresponds to a reassortant virus.

To classify a sample as coinfected based on one of the six internal gene segments (PB2, PB1, PA, NP, M, and NS), a different approach was needed because the internal segments of lineages I and II share 94-98% average nucleotide identity. As such, most internal gene segment reads would map to the reference genomes for both lineage I and lineage II, and coinfection could therefore be concluded erroneously. To assess coinfection at these internal gene segments, we identified the set of single nucleotide polymorphisms (SNPs) that differentiated lineage I from lineage II in each of the six internal gene segments (**Table S3**). Each of these differentiating SNPs (dSNPs) was characteristic of and unique to one of the two lineages. For each sample, we counted the number of lineage I and the number of lineage II dSNPs that were supported by the mapped reads (**Figure 3B**). Based on these results, we classified a given internal gene segment as either singly infected with lineage I, singly infected with lineage II, or coinfected (**Table S2**). Once every gene segment of a sample was classified as either singly infected or coinfected, we summarized these results at the sample level (**Figure 3C**, ‘All’). The majority of samples contained only lineage II gene segments (subtyped as H3N2). Samples containing only lineage I gene segments (subtyped as H1N1) were also apparent. We further found a non-negligible number of coinfected samples (*n* =70; **Table S2**), and a small number of reassortant samples (*n* = 9; **Table S2**).

To improve our ability to call variants in our downstream analyses, we selected 384 samples to re-sequence at even greater depth using two NovaSeq 6000 lanes (**Table S1, Table S2**). Re-sequencing of these samples resulted in a read depth of approximately 20,000 reads per nucleotide, substantially higher than the read depth in our initial sequencing runs (**Figure S1**).

### Within-host IAV diversity is low and largely synonymous in singly infected samples

To evaluate viral evolutionary dynamics without the complications introduced by co-infection, we decided to focus on within-host IAV diversity in pigs with samples that contained only lineage I or lineage II gene segments throughout their duration of infection. Samples classified as ‘unknown’ were presumed to derive from the same lineage as prior or later samples from the same pig and therefore did not affect categorization of pigs as persistently infected with lineage I or II. However, samples from pigs with any positive evidence of coinfection or reassortment were excluded. Among the re-sequenced samples, 260 from 168 pigs that met these criteria. We called intra-host single nucleotide variants (iSNVs) in these 260 samples using a variant calling threshold of 3% at sites that exceeded 500x coverage. The majority of these samples contained only low levels of IAV diversity, with 90% of the samples containing fewer than 15 iSNVs (**Figure 4A**). Most identified iSNVs were present at low frequencies of <10% (**Figure 4B**), paralleling findings in human IAV infections [19,20]. Overall, for both lineage I and lineage II samples, nonsynonymous iSNVs were present at lower frequencies than synonymous iSNVs, consistent with patterns one would expect in the presence of purifying selection. We further found that the ratio of the number of nonsynonymous to synonymous iSNVs generally fell below the neutral expectation, given by the ratio of nonsynonymous to synonymous sites (**Figure 4C**). This was the case for both lineages and across genes, and again points towards the role of purifying selection in shaping IAV diversity within swine. In two instances (lineage I M gene and lineage I NS gene) the calculated NS/S ratio fell above the neutral expectation but in these cases, the 95% confidence intervals were exceptionally broad, with lower bounds that fell below the neutral expectation.

**Figure 4.**
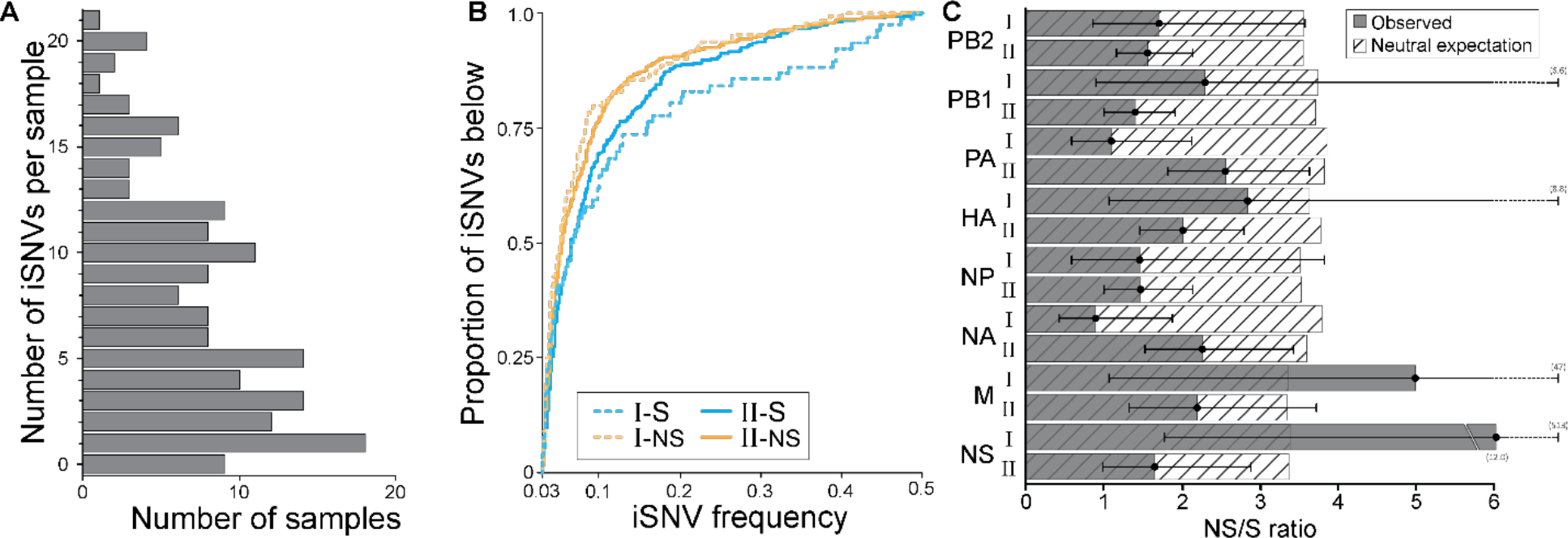
Singly infected samples harbor relatively low levels of genetic diversity and show evidence of purifying selection. **(A)** Distribution of the number of iSNVs identified per sample, across samples. (B) The proportion of detected iSNVs that fall below a given frequency, as specified on the x-axis. Approximately 70% iSNVs are detected at frequencies below 10%. Results are shown by sample lineage (I or II) and separately for nonsynonymous and synonymous iSNVs. (C) The ratio of the number of nonsynonymous to synonymous iSNVs, by gene segment and lineage. These ratios are shown alongside the neutral expectation, given by the ratio of nonsynonymous to synonymous sites. Black whiskers show the 95% confidence interval of the ratio.

The presence of spurious iSNVs in our dataset would result in NS/S ratios that are closer to the neutral expectation than might otherwise be the case. Because of this, we plotted the number of iSNVs called in a sample against the Ct value of the sample (**Figure S2**). A positive association between Ct value and the number of iSNVs could indicate that PCR amplification during sequencing may have generated spurious iSNVs. Despite our conservative variant calling threshold and our high coverage requirement, we did observe a positive association between Ct value and the number of iSNVs, indicating that some of the iSNVs that we detected are likely spurious. As such, the observation that nonsynonymous to synonymous ratios generally fall below the neutral expectation despite the likely presence of spurious iSNVs reinforces our conclusion that purifying selection is active. Finally, the presence of spurious iSNVs in our dataset would contribute to the number of iSNVs observed, and as such, levels of IAV diversity are likely lower than those shown in **Figure 4A**.

### A strong role for genetic drift in shaping within-host IAV evolution

To further characterize the evolutionary processes acting on swine IAV populations within singly infected pigs, we analyzed patterns of viral diversity in pigs with two or more longitudinal samples available. **Figure S3** shows iSNV dynamics, by segment, for all singly infected pigs with at least two resequenced time points (*n* = 82 pigs). To summarize these dynamics, we plotted the number of iSNVs in a sample against the timepoint during the pig’s infection at which the sample was taken (**Figure 5A**), as given by the number of days following the pig’s first PCR positive sample. We did not find a temporal pattern in iSNV richness over the course of infection, mirroring the finding in human IAV infections, where the number of iSNVs does not appear to change over the course of infection [19].

**Figure 5.**
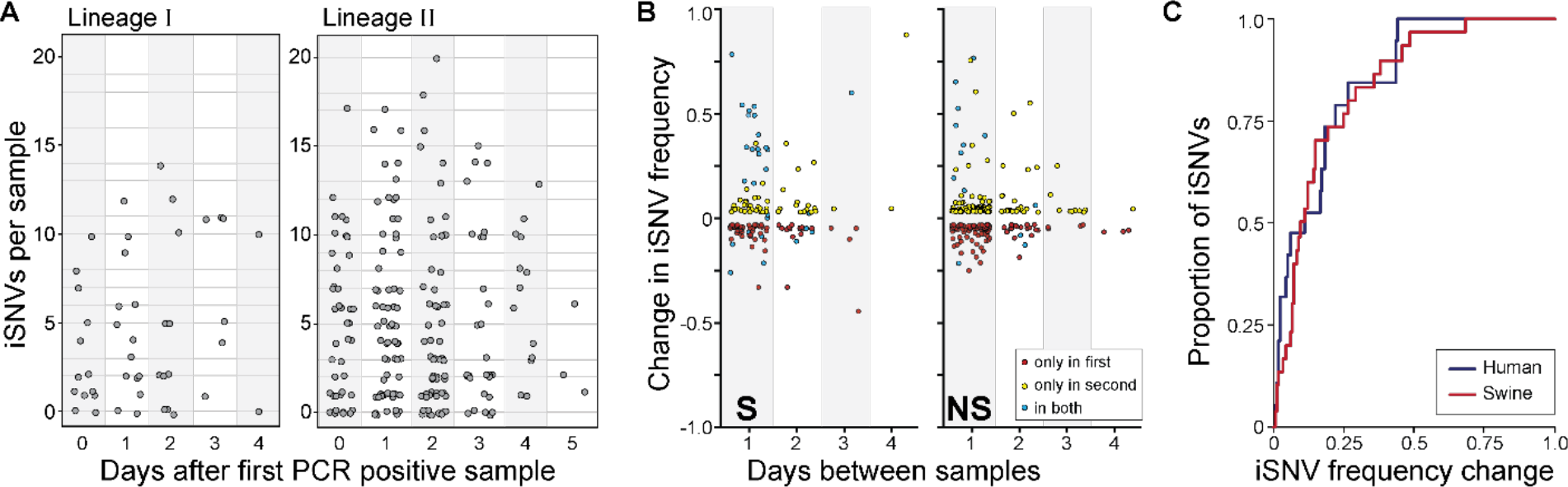
Longitudinal dynamics of iSNVs in singly infected swine. **(A)** The number of iSNVs detected in each sample according to the number of days since that animal had its first PCR positive sample, stratified by IAV lineage. (B) Changes in the frequency of synonymous (left), and non-synonymous (right) iSNVs according to the number of days between samples. iSNVs that were detected at both time points are shown in blue. (C) Cumulative distribution showing the proportion of iSNVs that persist between one day and the next with a frequency change that is less than or equal to the frequency change shown on the x-axis. Red line shows results from the swine data; the blue line shows results from humans IAV infections, as calculated from the data provided in [19]. For both datasets, we called iSNVs as present if found at frequencies of ≥3% and ≤97%.

To further quantify patterns of within-host viral evolution, we calculated, as in [19], changes in iSNV frequencies from one sequenced sample to the next sequenced sample. We plotted these changes as a function of the time between sample collection, stratifying by whether the iSNV was a nonsynonymous or a synonymous variant (**Figure 5B**). The apparent similarity in pattern between the nonsynonymous and synonymous iSNV frequency changes points towards the role of genetic drift playing a large role in within-pig IAV evolution. Considering all iSNVs together, we further found that a large proportion of iSNVs are only observed once in a pair of samples sequenced one day apart (**Figure 5B**). Specifically, in the subset of cases where there was sufficient depth and sequence quality in the later time-point to make an iSNV theoretically detectible, we calculated that 34% of iSNVs identified on the first day were not observed on the following day. Similarly, 30% of iSNVs identified in the second sample of a pair were not observed in the sample taken one day earlier. In comparison, using the data provided in [19] with an analogous variant-calling threshold of 3%, we calculated the percentage of iSNV loss and gain in human IAV infections to be 24% and 50%, respectively, between samples taken a day apart. In both swine and humans, the frequencies of iSNVs that persisted between samples taken one day apart often times changed dramatically (**Figure 5B**), with >50% of iSNVs that persisted from one day to the next exhibiting greater than a 12% change in frequency (**Figure 5C**). Further, the frequency change patterns in our swine data are remarkably similar to those calculated from the human IAV data provided in [19], suggesting that the strength of genetic drift acting on IAV populations is similar in pigs and humans.

## Discussion

Transmission of IAV among swine occurs commonly at agricultural fairs, where animals often raised on small farms from geographically disparate locations come together for a week long public event. Here, dense sampling of one such outbreak yielded a valuable set of IAV-positive samples representing both H1N1 and H3N2 subtype viruses and including time series that allow longitudinal evaluation of within-host viral dynamics. Several of the features of IAV populations in swine that we observed parallel those previously described in humans. Most notably, in both hosts, relatively few iSNVs are detected within a given individual and their frequency is typically low, indicating that mutation does not yield high diversity over the course of an acute infection [16,17,19,21]. Detected variants are furthermore often transient: similar to the patterns seen here in pigs, in humans, many iSNVs are not maintained above the limit of detection between longitudinal samples [19]. When variants are detected across adjacent samples, their frequencies can change dramatically, indicative of high levels of genetic drift. Purifying selection also appears to play a role in swine infections, as they do in human infections [17,19,20]. Interestingly, a recent study focused in young children revealed caveats to this trend in the case of infection with (antigenically novel) pandemic H1N1 virus [21]. In children infected with seasonal H3N2 viruses, the overall NS/S ratio fell below the neutral expectation (as seen in adults), but rates of nonsynonymous evolution increased over time after symptom onset. For children infected with pandemic H1N1 viruses, NS/S ratios instead fell above the neutral expectation, indicative of positive selection. IAV populations in pigs thus more closely mirror those in humans likely to have pre-existing immunity rather than those in children infected with pandemic H1N1 IAV. This might be because many swine included in our study are likely to have been vaccinated against influenza or because they may have had prior exposure to IAV.

While the dynamics of IAV populations in naturally infected humans have been examined in detail, studies to date in non-human hosts largely focus on experimental infections. Nonetheless, commonalities are apparent. In horses, dogs, pigs and quail infected experimentally with IAV strains that circulate naturally in these species, within-host nucleotide diversity was similarly low to that seen here in naturally infected pigs and in humans [22–25]. Rapid turnover of iSNVs has also been reported for multiple hosts, suggesting that the role of genetic drift in shaping within-host IAV populations is common to diverse host species [23–25]. Although the use of differing experimental and analytical approaches impedes direct comparison of NS/S ratios across hosts, available evidence in most non-human hosts examined suggests purifying selection is present as in adult humans and, as we found here, pigs.

The low viral diversity and predominance of genetic drift observed here in swine and generally in IAV infected hosts has important implications for viral evolutionary potential. As genetic diversity is a prerequisite for evolution, low diversity within hosts is indicative of relatively low adaptive potential. In addition, populations subjected to high levels of genetic drift are likely to adapt more slowly, as chance events can lead to the spread of deleterious mutations and loss of beneficial mutations. Stochastic effects therefore offer relevant explanations for the rare detection of antigenic variants within individual hosts, including those with prior immunity [16,19]. Notably, patterns of IAV evolution within acutely infected hosts stand in contrast to those observed at the level of host populations. In the global human population, IAV evolution is characterized by recurring selective sweeps of antigenically novel variant viruses [26–28]. Antigenic evolution is also apparent at a population level in swine [8]. However, a greater level of IAV diversity is maintained in swine populations compared to humans, with multiple antigenically distinct lineages co-circulating in a given geographical area [8]. This pattern is consistent with lower fitness advantages of new antigenic variants in swine populations relative to those in human populations, which could be explained by lower levels of host immunity in swine populations due to their shorter lifespans. Positive selection of antigenic variants being readily observed at the population-level, but not at the within-host level, argue for either only a small subset of infected individuals driving viral adaptation at the population-level or selection occurring at the between-host rather than at the within-host scale [29].

Our study has limitations that are important to consider in interpreting the data. Viral samples were collected with nasal wipes and the extent to which they are representative of the full viral population within an animal is unclear. Recent work showing strong spatial structure of IAV populations within mammals suggests that diversity present in the lower respiratory tract would not be efficiently sampled by this method [30,31]. In addition, although H1N1 and H3N2 subtype IAVs co-circulated in the sampled swine population, we excluded heterosubtypically coinfected animals from our analyses, thereby excluding viral diversity that would be generated through heterosubtypic coinfection and subsequent reassortment. This exclusion was made with the goal of focusing our analyses on a cohesive set of biological processes but, in the field, coinfection is likely to be an important source of IAV genetic diversity in swine. Conversely, our analysis likely includes some homosubtypically coinfected animals that were not defined as such; if present, such coinfected samples might inflate within-host genetic diversity. This potential concern is however mitigated by the observation that the number of detected iSNVs in our samples was low overall. Finally, all efforts to examine viral population diversity are subject to limitations on our ability to detect minor variants below a certain threshold. Subject to these limitations, our data indicate that IAV populations infecting farmed swine are characterized by limited genetic diversity and largely shaped by purifying selection and genetic drift similar to IAV populations within human hosts.

## Materials and Methods

### Ethics statement

Sampling of swine was reviewed and approved by The Ohio State University Institutional Animal Care and Use Committee under protocol number 2009A0134.

### Sample collection

Nasal wipes [32] were collected upon arrival at the agricultural fair during the swine exhibition weigh-in procedure, and then subsequently every night of the fair at 24-hour intervals as previously described in detail [33]. In brief, individual pig identification tags were recorded with samples, which were preserved on dry ice in the field after collection and for transportation to laboratory and long-term storage at -80°C. IAV infection was assessed using rRT-PCR targeting the M segment: National Veterinary Services Laboratory PCR primer protocol (no. SOP-BPA-9034.04) with SuperScript One-Step RT-PCR (Invitrogen). Samples were defined as IAV positive using a Ct cutoff of <45.0. Virus isolation on MDCK cells was attempted on a subset of samples and HA/NA subtypes of the viral isolates were evaluated using subtyping rRT-PCR (VetMAX™-Gold SIV Subtyping Kit; Life Technologies, Austin, TX USA).

### RNA extraction and sequencing

#### Initial Sequencing

All samples from days 1 (after the initial weigh-in procedure) through 6 of the fair that had Ct values of ≤31.0, based on M gene rRT-PCR, were initially sequenced. We refer to sequence data from this initial round of sequencing as Run 1. Influenza viral RNA extraction was performed using a custom QIAamp 96 DNA QIAcube treated kit (Qiagen) with a high-throughput automated liquid handler-QIAcube HT (Qiagen). Amplicons for sequencing libraries were generated through multi-segment RT-PCR (MRT-PCR) [34]. The resulting amplicons were then quantified using Quant-iT dsDNA High Sensitivity Assay (Invitrogen) and assessed by QIAxcel Advanced System (Qiagen) for size confirmation and presence of amplicon segments. The Nextera XT Sample Prep kit (Illumina) and Nextera XT Index kit v2 (Illumina) were used to produce paired-end DNA libraries using half-volume reactions. The amplicon libraries were purified using 0.8× AMPure XP beads (Beckman Coulter Inc.) on a Zephyr Compact Liquid handling workstation (Perkin Elmer). Purified libraries were then normalized and pooled using High sensitivity Quant-iT dsDNA to assess concentration and mean library size assessed by the QIAxcel. Six pM of the pooled libraries, including 5% PhiX, was loaded into a MiSeq v2 300 cycle kit (2x150bp) and MiSeq (Illumina) sequencer.

#### Resequencing

We selected samples for resequencing using the following priorities: (1) For every singly infected pig with only a single sequenced sample, we included this sample for resequencing. (2) For every singly infected pig with 2 or more longitudinal samples, we selected the earliest sample and the latest sample for resequencing. This resulted in 110 pairs of longitudinal samples taken 1 day apart, 48 pairs of samples taken 2 days apart, and 27 pairs of samples taken 3 or more days apart. (3) If any sample chosen in (2) had low sequence coverage in the first run and a high-Ct matrix rRT-PCR value, that sample was replaced with an alternative sample from the same pig when possible. We refer to resequenced samples as Run 2 samples. RNA was re-extracted from frozen samples and libraries were prepared as described above, then 225pM of pooled library were loaded in the Illumina NovaSeq 6000 SP reagent kit v1.5 300 cycle (2x150bp) paired-end sequencing kits.

### Short read processing and alignment

Raw reads were processed using bbtools to remove sequencing adapters and trim low-quality bases [35]. We then used IRMA (Iterative Refinement Meta-Assembler) [36] to generate consensus sequence for each sample. These consensus sequences indicated that there were two IAV subtypes co-circulating at the fair: one from a swine H1N1 lineage and one from a swine H3N2 lineage (referred to as lineages I and II in this manuscript, respectively). For any given segment, the within-lineage variation in consensus sequence was low (average pairwise distance was within one nucleotide difference). Accordingly, the major haplotype of each segment from each lineage was selected to be used as a reference sequence (16 total reference sequences: 8 from lineage I and 8 from lineage II). Reads from each sample were split according to segment and subtype to which they shared the greatest nucleotide similarity. To do this, all reads were aligned to the 16 reference sequences using BLAT [37]. Each subtype and segment specific bin of short reads was then mapped onto their respective reference sequence using bbmap with the following settings: 90% minimum percent identity, toss ambiguously mapped reads, and global alignment mode. The coverage for all samples, by segment, is shown in Figure S4.

### Reconstruction of haplotype networks and identification of differentiating SNPs (dSNPs)

Using the Run 1 sequencing data, consensus sequences were generated for all segments in each sample by identifying the major allele at each site with ≥10x coverage. Sites with <10x coverage were recorded as gaps. Since the internal gene segments (PB2, PB1, PA, NP, M and NS) share on average 93-97% nucleotide identity between lineages, reads were mapped separately to lineage I and lineage II references, but total coverage and allele frequencies were calculated using alignments to both references. This method was adopted because, although the similarity of internal segments was sufficiently high to permit mapping of all reads to the same reference, this practice made it difficult to evaluate the significance of low-frequency variants. Low quality samples and potentially coinfected samples were excluded from alignments by removing all samples that contained ≥1% polymorphic sites. Using these alignments containing only sequences with high coverage from samples unlikely to be coinfected, we inferred haplotype networks using a custom python script (GitHub: https://github.com/Lowen-Lab/swineIAV). SNPs that differentiate lineage I from lineage II viruses (dSNPs) were identified as those SNPs that distinguish the dominant lineage I and lineage II haplotypes. The number of detected dSNPs for all samples is shown in **Figure S4**.

### Classifying sequenced segments and samples into lineage designations

Because lineage I and II HA and NA segments share little nucleotide similarity, short reads map reliably to only one reference or the other. Mapping coverage is therefore a reliable method of distinguishing H1 from H3, and N1 from N2. Accordingly, for HA and NA segments, positive evidence of lineage I or II genotypes required that reads map to at least 10% of all sites in a segment with 50x coverage or higher in Run 1 and 500x coverage in Run 2.

Internal segments (PB2, PB1, PA, NP, M and NS) were classified into lineages based on the alleles present at the dSNPs sites. The base quality and mapping quality thresholds to count a putative dSNP allele were set empirically based on the distribution of quality and mapping statistics observed in our sequence data and using our read alignment protocol. Specifically, only sites with ≥50x coverage (or 500x for re-sequenced samples), an average mapping score of the major allele ≥43, and average mismatch and indel counts of reads containing the major allele ≤1.5 and ≤0.5, respectively. At those sites, only those alleles present at ≥3% frequency, with an average phred score of ≥37, average mapping quality score of ≥40, and average location of that allele in each mapped read ≥30 bases from the nearest end of a read were included. A sample’s gene segment was classified as supporting infection with a lineage I gene segment if more than 10% of the total dSNP sites for a segment contained the alleles defining lineage I. Similarly, a sample’s gene segment was classified as supporting infection with a lineage II gene segment if more than 10% of the total dSNP sites for a segment contained the alleles defining lineage II. If a sample’s gene segment supported infection with both a lineage I and a lineage II gene segment, it was classified as coinfected. Gene segments that were not classified as either lineage I or lineage II were classified as “unknown”. As such, gene segments with low sequencing coverage were generally classified as “unknown”.

At the sample level, a sample was classified as coinfected if at least one of the eight segments was classified as coinfected. Of the remaining samples, those with five or more segments classified as lineage I or lineage II were classified as lineage I or II, respectively, at the sample level. Samples with five or more segments classified as “unknown” were classified as “unknown”. Samples with some segments classified as lineage I and others classified as lineage II were classified as reassortant at the sample level, but only if the number of “unknown” segments was less than five. We manually inspected the classification of all samples to verify the accuracy and consistency of the genotyping process.

### Identification of within host single nucleotide variants (iSNVs)

Run 2 data were used for calling iSNVs. Similar to the identification of dSNPs, the cutoffs for including iSNVs were set empirically. First, sites were evaluated based on their total coverage and the average quality and mapping statistics of reads containing the major allele. In particular, only sites with ≥500x coverage, an average mapping score of the major allele ≥43, and average mismatch and indel counts of reads containing the major allele ≤1.5 and ≤0.5, respectively, were included. These major allele focused site exclusion criteria were introduced out of necessity and functioned primarily to exclude false positives driven by mapping errors, such as at the borders of defective viral genomes (DVGs). For minor variants at these sites to be included in subsequent analyses, they were required to be present at ≥3% frequency, have an average phred score of ≥37, and the reads that contained the minor allele at any given site also had to have sufficient mapping quality to justify inclusion. Specifically, reads containing the minor allele needed an average mapping quality score of ≥40, the average location of the minor allele needed to be ≥30 bases from the nearest end of a read, and the reads overall needed to have ≤2.0 and ≤1.0 average mismatch and indel counts, respectively, relative to the reference sequence.

## Acknowledgments/Funding

This study was supported by National Institute of Allergy and Infectious Diseases (NIAID) Centers of Excellence for Influenza Research and Response (CEIRR) contract no. 75N93021C00017 and an Emory University MP3 seed grant. Further support for this study was provided by the US National Institutes of Health National Institute of General Medical Sciences grant 1R01 GM124280-03S1 (supplement). Funding for open access charge: US National Institutes of Health National Institute of General Medical Sciences grant (1R01 GM124280-03S1). Ohio State’s contributions were supported by CEIRR contract 75N93021C00016 and the Centers for Disease Control and Prevention through cooperative agreement U38OT000143 with the Council of State and Territorial Epidemiologists. The findings and conclusions in this report are those of the authors and do not necessarily represent the views of the Centers for Disease Control and Prevention or the Agency for Toxic Substances and Disease Registry.

## Supplementary information captions

### Supplementary Tables

**Table S1**. Nasal wipes collected over the week-long county fair. Columns include vial numbers, pig identification numbers, sampling day, Ct value from the rRT-PCR assay targeting the M segment, infection status based on this assay, and subtype-specific assay results, when performed. A sample was classified as PCR positive for IAV if the Ct value of the qRT-PCR assay targeting the M segment was ≤45.0. Sequence identification numbers of samples that were sequenced are also provided. These unique sequence IDs correspond with those deposited to the NCBI SRA ([pending CDC review, insert bioproject number]).

**Table S2**. Table of sequenced samples. The first 8 columns of the table include the vial number, pig ID, sampling day, Ct value from the rRT-PCR assay targeting the M segment, and subtype-specific assay results (when available). The next 10 columns of the table include the unique sequence ID from Run 1, the classification of each gene segment into a lineage based on the deep sequencing data, and the classification of the sample into a lineage based on all viral gene segments. The last 10 columns include analogous information and results based on Run 2, when applicable.

**Table S3**. List of differentiating SNPs (dSNPs) for the six internal gene segments. Each tab lists the gene segment, the dSNP nucleotide site, and the nucleotides present at the corresponding dSNP sites for lineage I and lineage II viruses.

## Supplementary Figures

**Figure S1.**
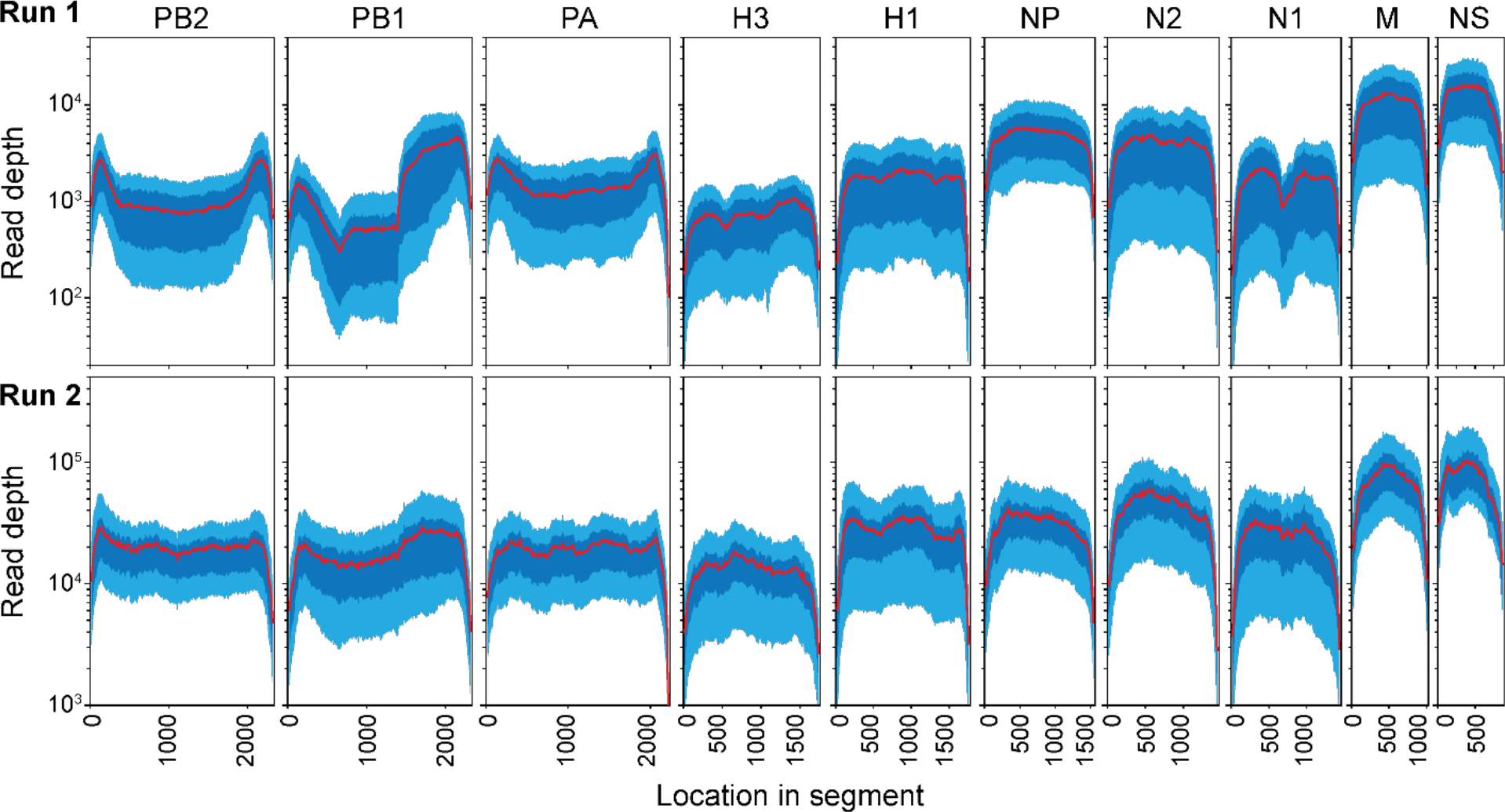
Coverage for sequenced samples. The x-axis shows the location along the IAV genome. The y-axis shows read depth. Coverage for Run 1 and Run 2 samples are shown separately. A sliding window of 200 nucleotide sites was used to calculate mean read depth for each sample at each site. The plotted ranges show median (red), 25^th^ (dark blue), and 75^th^ (light blue) percentiles of read depth across all samples sequenced in Runs 1 and 2.

**Figure S2.**
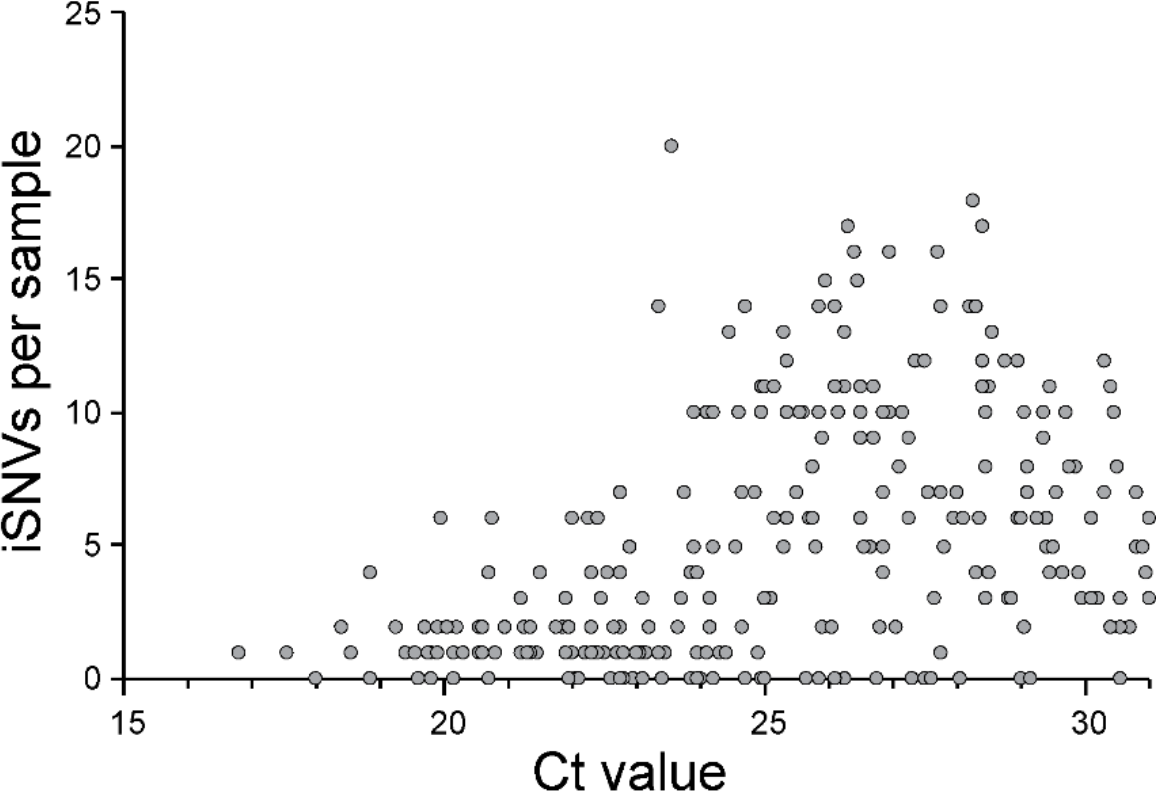
Scatterplot showing the relationship between the rRT-PCR Ct value of a sample and the number of iSNVs detected in the sample. A positive relationship is evident, with higher Ct samples more likely to have a larger number of iSNVs detected.

**Figure S3.**
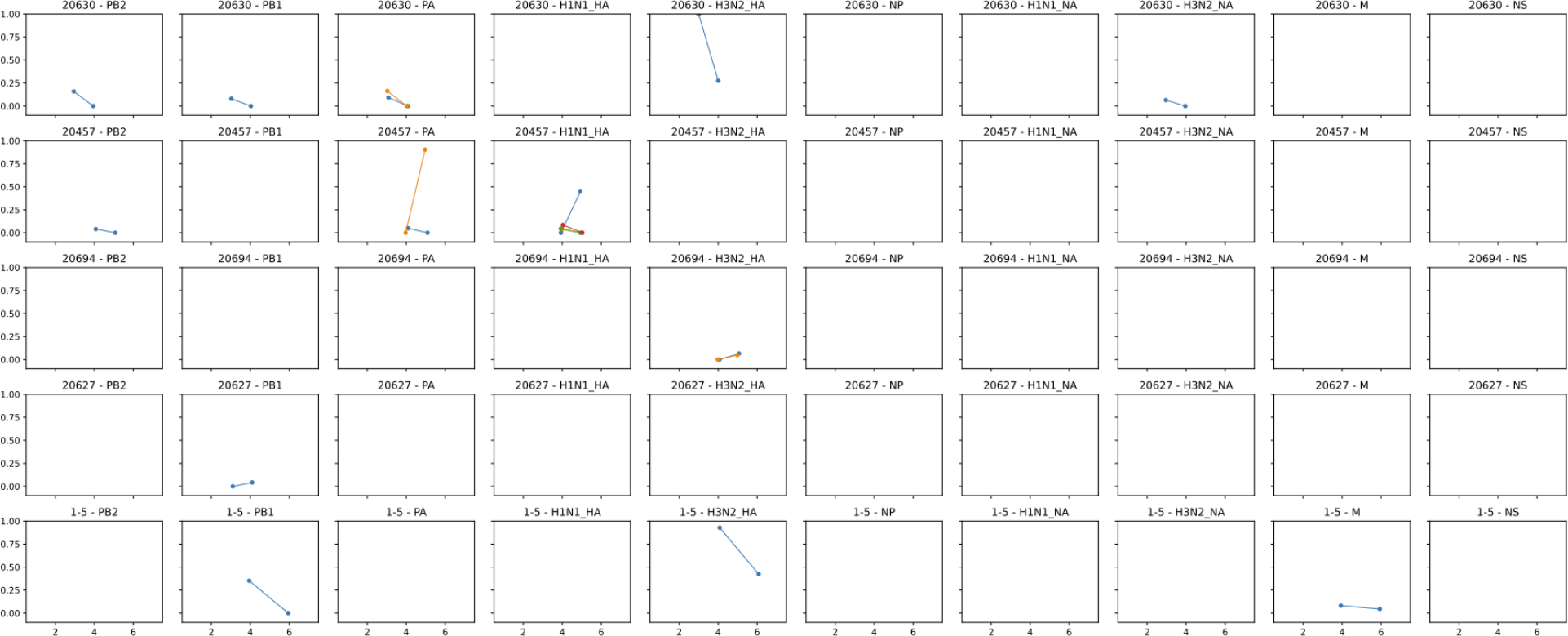
iSNV dynamics within singly infected pigs with two or more sequenced samples. Each row shows iSNV dynamics for a single pig. Columns correspond to the six internal gene segments of IAV, and both lineage I and II HA and NA gene segments. Blank graphs indicate no iSNVs were detected in the reads mapping to each reference sequence. The x-axis specifies the sample day and the y-axis specifies iSNV frequency. ***(Figure above is representative only, see attached PDF for full length figure)***.

**Figure S4.**
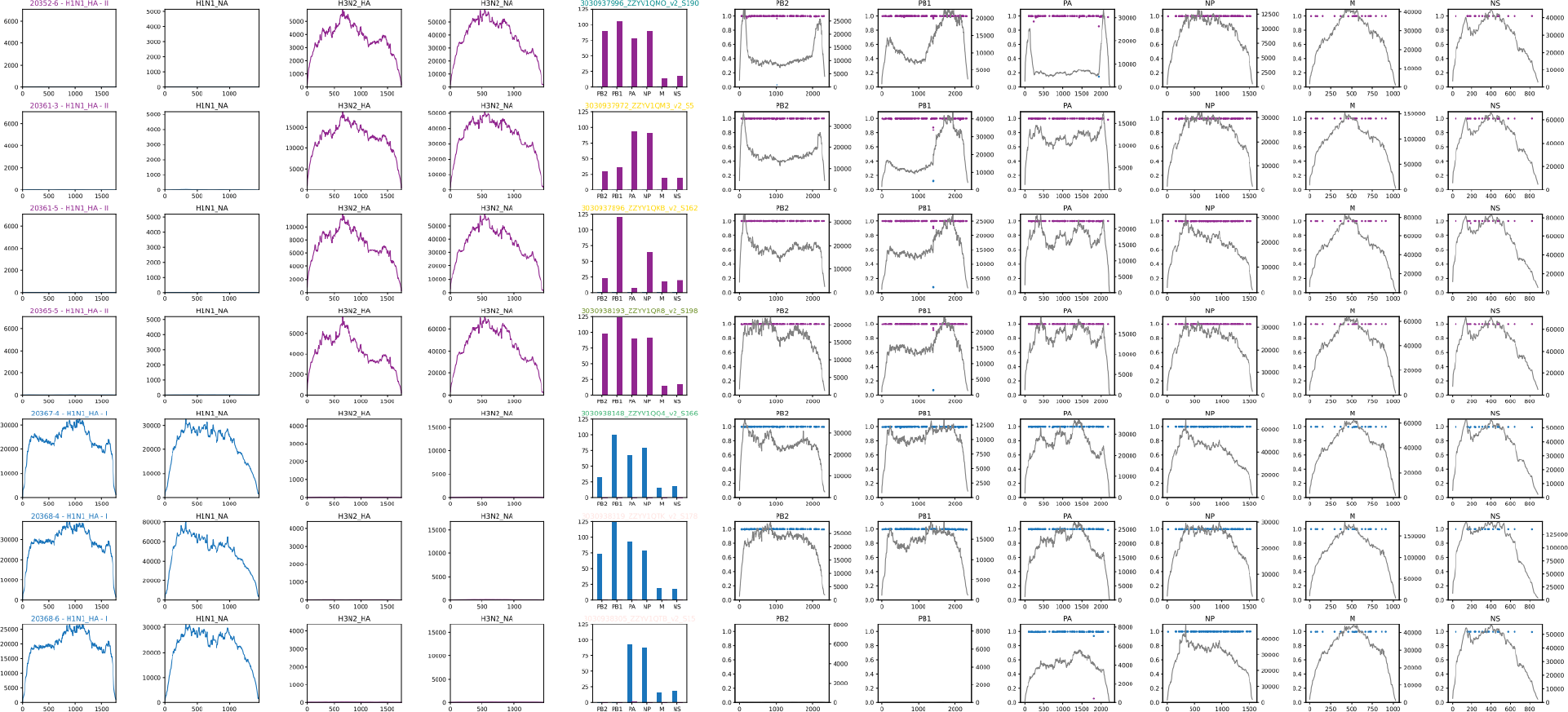
Coverage and the number of detected dSNPs for all samples, by segment. Each row is a separate sample. The pig ID, sample day, and sample classification are shown in the panel titles of the fifth column. The first four columns show mapping coverage against H1, N1, H3, and N2 references, respectively. The fifth column shows the number of dSNPs detected for each lineage in each of the six internal segments. The remaining six columns show coverage and dSNP frequency by location in the internal segments in the following order: PB2, PB1, PA, NP, M, NS. ***(Figure above is representative only, see attached PDF for full length figure)***.

